# PAM: Predictive attention mechanism for neural decoding of visual perception

**DOI:** 10.1101/2024.06.04.596589

**Authors:** Thirza Dado, Lynn Le, Marcel van Gerven, Yağmur Güçlütürk, Umut Güçlü

## Abstract

In neural decoding, reconstruction seeks to create a literal image from information in brain activity, typically achieved by mapping neural responses to a latent representation of a generative model. A key challenge in this process is understanding how information is processed across visual areas to effectively integrate their neural signals. This requires an attention mechanism that selectively focuses on neural inputs based on their relevance to the task of reconstruction — something conventional attention models, which capture only input-input relationships, cannot achieve. To address this, we introduce predictive attention mechanisms (PAMs), a novel approach that learns task-driven “output queries” during training to focus on the neural responses most relevant for predicting the latents underlying perceived images, effectively allocating attention across brain areas. We validate PAM with two datasets: (i) B2G, which contains GAN-synthesized images, their original latents and multiunit activity data; (ii) Shen-19, which includes real photographs, their inverted latents and functional magnetic resonance imaging data. Beyond achieving state-of-theart reconstructions, PAM offers a key interpretative advantage through the availability of (i) attention weights, revealing how the model’s focus was distributed across visual areas for the task of latent prediction, and (ii) values, capturing the stimulus information decoded from each area.

## Introduction

Attention mechanisms in deep learning draw inspiration from the cognitive ability to selectively focus on specific aspects of the environment while neglecting others (Kastner & Ungerleider, 2000). These computational models dynamically weigh the importance of different input data segments to prioritize the most relevant information for the task at hand (Bahdanau et al., 2014; Vaswani et al., 2017) – just as humans focus their attention on key details to understand a scene or address a problem. In brief, an attention model derives three components from the input data: queries, keys and values. A query acts like a spotlight, shaped by a specific objective, to identify which parts of the input data are most pertinent (e.g., in a language translation model, the query could be the representation of a word in a sentence for which the model seeks the best equivalent in the target language). Keys are representations of the input data that embed contextual information (e.g., in a language translation model, a key associated with a particular word would also capture aspects of the surrounding words) so that the model can understand how each data segment fits into the larger picture. Keys are designed to be matched against queries to evaluate their relevance, and their compatibility results in the corresponding attention weight. Values carry the actual output information and are aggregated according to the attention weights (e.g., in a language translation model, values could be potential translations for words or phrases). Through this mechanism, a model dynamically prioritizes the most relevant parts of the input by calculating an attention-weighted sum of the values.

Neural decoding addresses the inverse problem of translating neural activity back into the features of a perceived stimulus to which the brain responds (Fig 1). This process seeks to find how the characteristics of a phenomenon are represented in the brain through classification (Haxby et al., 2001; Kamitani & Tong, 2005; Stansbury et al., 2013; Huth et al., 2016; Horikawa & Kamitani, 2017), identification (Mitchell et al., 2008; Kay et al., 2008; Gü çlü & van Gerven, 2017a,b) or reconstruction (Thirion et al., 2006; Miyawaki et al., 2008; Naselaris et al., 2009; van Gerven et al., 2010; Nishimoto et al., 2011; Schoenmakers et al., 2013; Gü çlü & van Gerven, 2013; Cowen et al., 2014; Du et al., 2017; Gü çlü tü rk et al., 2017; Shen et al., 2019; VanRullen & Reddy, 2019; Dado et al., 2022, 2024). Among these, reconstruction is the most challenging, as it requires generating a literal replica of the perceived stimulus from brain activity alone. For visual perception, this can be achieved with state-of-the-art quality using a decoding method that uses generative adversarial networks (GANs) (Goodfellow et al., 2014) to synthesize visual information represented in the brain, as shown in previous research (Dado et al., 2022, 2024). In brief, a decoder is trained to map neural responses to GAN latent vectors (Fig 1: “decoding”), which are then fed to the GAN’s generator to reconstruct the corresponding images (Fig 1: “synthesis”). The latent vectors of the training set can be obtained in two ways: (i) a priori, by using images already generated by the GAN, where the latents are known in advance, or (ii) post-hoc, by retrospectively estimating latent vectors for given stimuli. In this work, we employed both approaches: for stimuli that originated from a GAN, we used their known latent representations (i); for real images, we implemented (ii) by optimizing a latent vector to generate an image that closely matches the training stimulus in terms of perceptual similarity.

**Figure 1:**
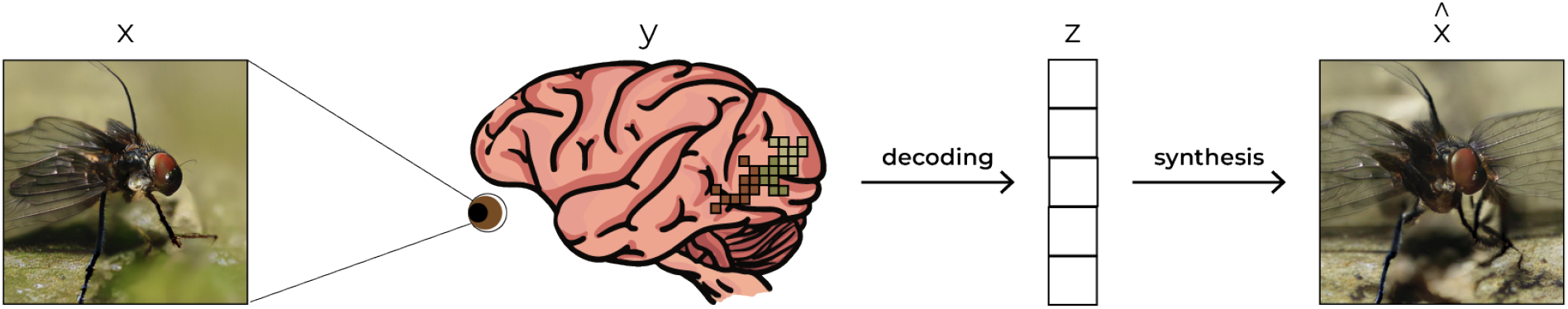
Neural decoding. This inverse problem seeks to infer the stimulus underlying the observed neural activity. It is common to divide this process into two stages: a “decoding” transformation that maps neural responses to an intermediate feature representation, and a more complex “synthesis” transformation that converts these features into an actual image.

A key challenge in reconstruction is to determine how different visual areas represent a perceived stimulus to effectively extract and combine this information. This calls for an attention mechanism that selectively focuses on neural inputs based on their relevance to the task of reconstruction. However, conventional attention models are not designed to establish relationships between neural inputs and output features (in our case, GAN latents) for reconstruction. For instance, self-attention, a widely used mechanism in Transformers, computes relationships between elements within the input itself (e.g., words within a sentence) such that keys and queries are both derived directly from the input. Once again, queries represent **what to look for**, while keys represent **what is available**. Attention weights are computed by comparing keys against queries, where higher matches indicate greater relevance. In our case, we seek to prioritize neural responses that are relevant for predicting (anticipated) feature outputs, rather than the relationships among the neural areas themselves. As such, we need goal-oriented “output queries” that guide attention toward neural signals relevant for predicting output features that do not yet exist at the time of computation.

To address this, we developed a predictive attention mechanism (PAM), which learned “output queries” as trainable parameters during training (Fig 2). Unlike input-derived queries, these output queries remain fixed after training and serve as a task-driven attention strategy, ensuring that attention is consistently directed toward the relevant neural signals for reconstruction. To evaluate PAM, we tested it on two datasets with distinct properties: one comprising intracranial multi-unit activity (MUA) recordings from macaque visual cortex with GAN-synthesized images and their original latents, and another using noninvasive functional magnetic resonance imaging (fMRI) data from human participants with real images and latents estimated retrospectively. Notably this work is the first to extend the GAN-based method of (Dado et al., 2024) to datasets of real images where latent representations were not available a priori, instead estimating them post-hoc.

**Figure 2:**
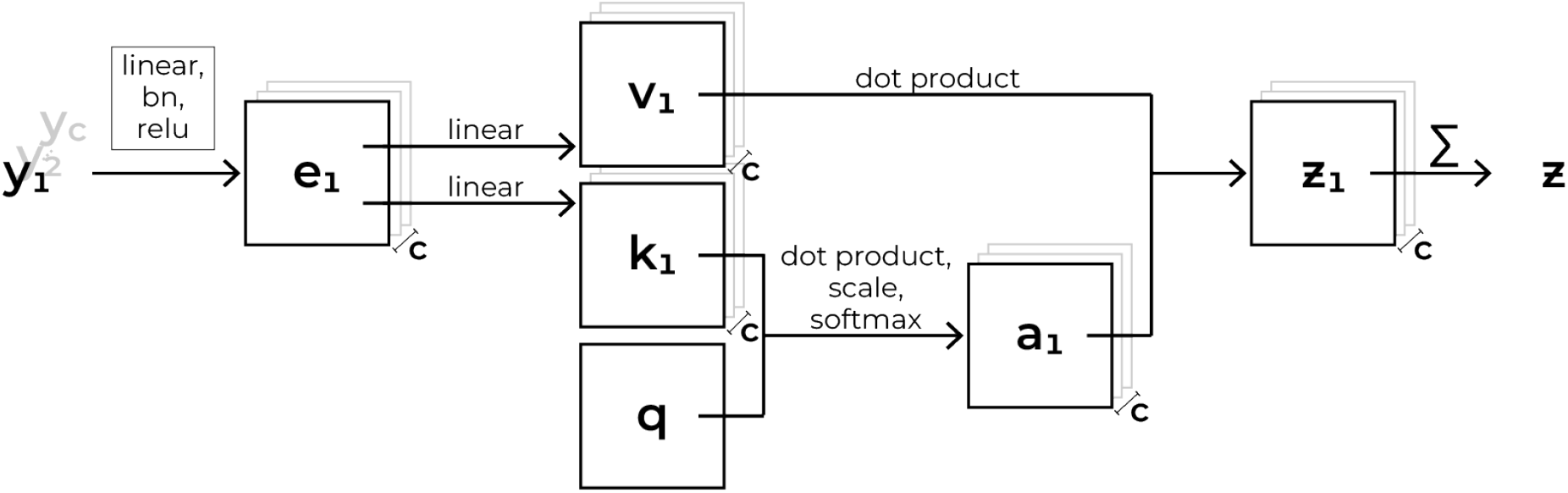
Predictive attention mechanism (PAM). The **input** data, *Y* = {*y*_1_, *y*_2_, …, y_c_} comprises neural data from *c* regions of interest (i.e., the number of attention heads) and the **output** *z* the decoded latent of the stimulus. First, *Y* is transformed via *n* nonlinear embedding blocks, where each block consists of a linear layer, batch normalization, and ReLU activation, producing an embedded representation *E* = {*e*_1_, *e*_2_, …, *e*_*c*_}. Keys *K* = {*k*_1_, *k*_2_, …, *k*_*c*_} and values *V* = {*v*_1_, *v*_2_, …, *v*} are then derived from *E* through a (separate) linear transformation, with each attention head having its own embedding, key and value transformations. Unlike *K* and *V*, the “output queries” *q* are learned during training and dynamically adapt to emphasize the most relevant neural features for the task of latent prediction. Output queries interact with the keys via matrix multiplication, scaling and a softmax operation to compute attention weights, *A* = {*a*_1_, *a*_2_, …, *a*_*c*_}. Finally, the attention-weighted sum of *V* yields the predicted output features (latent), *z*, which serve as the basis for reconstructing the stimulus.

## Methods

### Predictive attention mechanism

PAM (Fig 2) is specifically designed to determine the relevance of each neural area to predict the specific output features underlying a stimulus. The **input** data, *Y* = {*y*_1_, *y*_2_, …, *y*_*c*_}, comprises neural data from *c* regions of interest (i.e., the number of neural areas; the number of attention heads), and the **output** *z* represents the predicted output features (i.e., the GAN latent) underlying the stimulus, which is fed to the GAN to synthesize the corresponding image. We utilized StyleGAN-XL (Sauer et al., 2022), trained on ImageNet (Deng et al., 2009), to generate 512 × 512 pixel images from 512-dimensional feature-disentangled *w*-latents (rather than the *z*-latents). To ensure reconstructions faithfully reflected the full range of variability, we omitted both random noise and truncation, avoiding artificial bias toward highquality images.

First, the model embeds the neural input via multiple attention heads, where each head corresponds to the input from a specific brain area. This embedding applies nonlinear transformations to project neural responses into a feature space more suitable for decoding. Each input *y*_*i*_ undergoes *n* consecutive transformations, each consisting of a linear layer, batch normalization, and ReLU activation, yielding the embedded representation, *E* = {*e*_1_, *e*_2_, …, *e*_*c*_}. Formally, for each region of interest *i* (where *i* ∈ {1, 2, …, *c*}, indexing the *c* neural regions), this transformation can be represented as

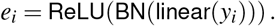

The number of embedding blocks was dataset-dependent: for Shen-19 (fMRI data), we used a single block, whereas for B2G (MUA data), we applied five blocks, allowing deeper nonlinear transformations. This choice was based on empirical results: additional blocks did not improve fMRI decoding but significantly enhanced MUA decoding, likely due to the higher signal quality of MUA data.

Next, keys *K* = {*k*_1_, *k*_2_, …, *k*_*c*_} and values *V* ={*v*_1_, *v*_2_, …, *v*_*c*_} are derived from *E* using linear transformations

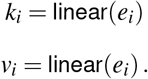

Each attention head has its own embedding, key, and value transformation. Unlike *K* and *V*, the queries *q* are not predefined or explicitly computed but learned during training. Specifically, they are updated along with the rest of the model’s parameters by minimizing the mean squared error (MSE) between the predicted and target latents (see training details below). This means that, over time, the queries self-organize to prioritize neural signals most informative for predicting the correct latents for reconstruction. The keys then interact with these learned queries to determine the focus of attention through matrix multiplication, scaling, and a softmax operation (i.e., the scaled dot product (Vaswani et al., 2017)) to compute attention weights, *A* = {*a*_1_, *a*_2_, …, *a*_*c*_}

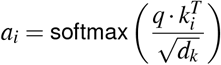

where *d*_*k*_ is the dimension of the key vectors. The division by 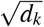 stabilizes attention computation, and the softmax function ensures that the attention weights sum to one.

The final output is a weighted sum of the value vectors and attention weights, allowing PAM to emphasize neural embeddings most relevant to the latents. This yields the predicted output features, z

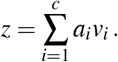

The model was trained to minimize the MSE loss between the predicted and target latents

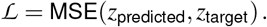

We optimized the model using Adam with default parameters and initialized the weights with Xavier uniform initialization to ensure stable gradient flow. Training was performed with a batch size of 32 and continued until convergence.

### Neural datasets

We used two datasets comprising naturalistic images and corresponding brain responses to decode perceived images from neural activity. The first dataset, **“B2G”**, consists of synthetic images generated by StyleGAN-XL with readily available latent vectors, providing a controlled setup for evaluating the decoding process. It contains invasive multi-unit activity (MUA) recordings — the envelope of the spiking activity measured using multi-electrode Utah arrays — from visual areas V1, V4 and IT in one macaque, as detailed in (Dado et al., 2024). In total, B2G consists of 4000 and 200 training (1 repetition) and test (20 repetitions) examples, respectively. The preprocessed MUA data is available at Figshare at DOI 25637856. The second dataset, **“Shen-19”**, comprises natural images from ImageNet paired with noninvasive fMRI data from seven visual areas (V1-V4, FFA, LOC, PPA) in three human participants, as detailed in (Shen et al., 2019). It contains 1200 training examples (five repetitions) and 50 test examples (24 repetitions). The preprocessed fMRI data is available at Figshare at DOI: 7033577/13.

#### fMRI preprocessing

We applied *hyperalignment* to the fMRI recordings *per brain area* in the Shen-19 dataset to map subject-specific responses into a shared functional space (Haxby et al., 2020). This procedure accounts for individual differences in brain anatomy and functional topology, allowing responses to be averaged across participants while preserving region-specific functional similarity. After transforming each participant’s data into this common space, we computed the mean response across the three participants, yielding a representative response pattern while reducing inter-subject variability. We z-scored the averaged training and test responses using statistics from the training set to ensure normalized feature scaling for subsequent analyses.

We *selected informative voxels* using ridge regression with 10-fold cross-validation to predict voxel responses from latent vectors of the training examples. To optimize regularization, we explored a range of lambda values for the ridge penalty, derived from the singular value decomposition (SVD) of the input matrix. Specifically, we extracted the nonzero singular values, using their square values to determine the upper_272_ and lower bounds for lambda selection. Five logarithmically spaced lambda values were generated between these bounds to ensure a well-distributed search space. Based on the Pearson correlation between the predicted and target responses_276_ on the training set, we selected voxels using a false discovery rate (FDR) corerction (α = 0.05) applied per visual area (see Table 1). This controls the expected proportion of “false discoveries” (erroneously rejected null hypotheses) in multiple hypothesis testing, ensuring that retained voxels exhibit statistically reliable predictive power.

**Table 1:** Voxel Selection. The number of voxels pre- and post-FDR thresholding, which is used to eliminate less reliable responses. The voxels that remain post-FDR are considered more likely to be truly responsive to the visual stimuli. Notably, the voxel count in the PPA was reduced to zero following FDR adjustment, suggesting a lower reliability in the initial responses from this area.

|  | before FDR | after FDR |
| --- | --- | --- |
| V1 | 2521 | 1643 |
| V2 | 2687 | 1860 |
| V3 | 2058 | 1451 |
| V4 | 1283 | 968 |
| LOC | 2724 | 2134 |
| FFA | 2100 | 1210 |
| PPA | 900 | 0 |
| Total | 14273 | 9266 |

For *latent inversion*, we optimized an input latent for each training image to best match the stimulus in VGG16 feature space, minimizing the learned perceptual image patch similarity (LPIPS) distance. To account for variability due to initialization, we repeated this process ten times with different random seeds and selected the latent with the lowest LPIPS distance, ensuring the closest perceptual match (see Fig 3).

**Figure 3:**
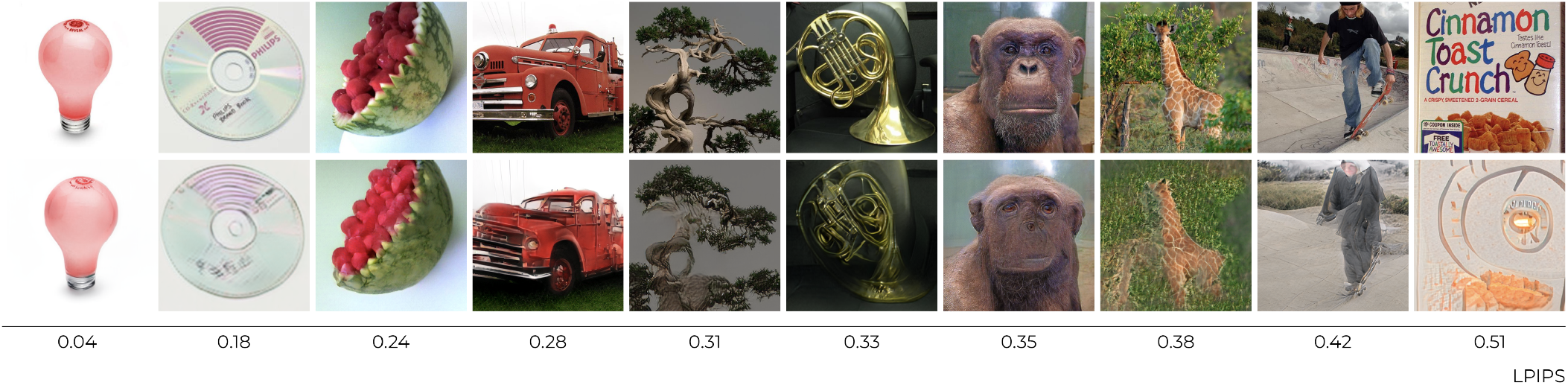
Latent inversion from real photographs. Ten arbitrary Shen-19 training stimuli (top) and their corresponding reconstructions from the inverted latents (bottom), with the LPIPS distance indicating dissimilarity between them. These examples span the full range of LPIPS, from high-similarity cases on the left (low LPIPS distance) to lower-similarity cases on the right (high LPIPS distance). Note that these latents were directly inverted from images, without decoding from brain activity.

### Evaluation

We quantified the alignment between original stimuli and their reconstructions using three cosine similarity-based metrics: learned perceptual image patch similarity (LPIPS), perceptual similarity, and latent similarity.

- LPIPS: Feature representations were extracted from multiple layers of VGG16, pretrained for object recognition.
- Perceptual Similarity: Feature representations were taken from five distinct VGG16 levels following max pooling, capturing different complexity levels. Lower layers reflected low-level image features, while higher layers represented more abstract characteristics.
- Latent Similarity: Cosine similarity between latent vectors of the original and reconstructed images.

### Implementation details

All analyses were conducted using Python 3.10.4 on a cloud-based virtual machine equipped with an AMD EPYC7F72 24-Core Processor (2.5 GHz 3.2 GHz) and 96 cores, running a Linux kernel version 4.18.0-372.80.1.el8 6 on an x86 64 architecture. We employed the original PyTorch implementation of StyleGAN-XL and used VGG16 for object recognition to measure perceptual similarity during evaluation. The code to reproduce the main experimental results can be found on our anonymous GitHub repository.

## Results

We trained two PAM-based decoders, each adapted to the specific characteristics of the B2G and Shen-19 datasets by varying the number of blocks in their embedding transformations. To assess PAM’s performance, we compared it to the linear GAN-based decoding baseline introduced by (Dado et al., 2024), which represents the state-of-the-art in neural reconstruction.

- **B2G (MUA data)**. The linear GAN-based decoder was originally developed for this dataset and serves as the established state-of-the-art baseline. In our experiments, PAM outperformed the baseline, benefiting from a five-block embedding transformation that captured complex mappings from high-resolution MUA signals to latent vectors. This resulted in improved reconstructions (Fig 4) and higher quantitative metrics (Table 2).
- **Shen-19 (fMRI data)**. GAN-based linear decoding had not been applied to this dataset before because Shen-19 contains real images without pre-existing latents. Here, we inverted latents for Shen-19 images, enabling the first application of the linear decoding method to a dataset of images not generated by a GAN and establishing a new baseline for comparison with PAM. Empirically, deeper embeddings did not improve decoding, so we used a single embedding block for this dataset. This is likely due to the higher noise levels in fMRI, its lower resolution and the smaller dataset size, which increase the risk of overfitting. The results show only marginal differences between PAM and the linear baseline, as both models extract similar features given PAM’s single-block embedding transformation.

**Table 2:** Reconstruction performance. Results (mean ± SE) for PAM (P) and the linear decoder (L) on B2G and Shen-19 datasets across metrics: LPIPS, perceptual similarity (VGG 2/16 to 13/16) and latent cosine similarity (Lat sim). Lat sim is unavailable for Shen-19 because the dataset uses real-world photographs without pre-defined latent vectors, unlike the GAN-synthesized images in B2G. *p*-values in the last row show significance from paired t-tests: for B2G, PAM outperforms linear decoding (*p* ≪ 0.0001); for Shen-19, differences are nonsignificant (*p* > 0.05).

|  |  | LPIPS sim | VGG 2 | VGG 4 | VGG 7 | VGG 10 | VGG 13 | Lat sim |
| --- | --- | --- | --- | --- | --- | --- | --- | --- |
| B2G | P | $0.36 \pm 0.007$ | $0.43 \pm 0.006$ | $0.35 \pm 0.004$ | $0.29 \pm 0.005$ | $0.28 \pm 0.009$ | $0.36 \pm 0.013$ | $0.84 \pm 0.006$ |
| | L | $0.32 \pm 0.005$ | $0.41 \pm 0.005$ | $0.33 \pm 0.004$ | $0.26 \pm 0.003$ | $0.22 \pm 0.004$ | $0.25 \pm 0.007$ | $0.77 \pm 0.003$ |
| | $p$ | $< 0.0001$ | $< 0.0001$ | $< 0.0001$ | $< 0.0001$ | $< 0.0001$ | $< 0.0001$ | $< 0.0001$ |
| Shen-19 | P | $0.26 \pm 0.009$ | $0.38 \pm 0.009$ | $0.30 \pm 0.006$ | $0.23 \pm 0.004$ | $0.18 \pm 0.005$ | $0.17 \pm 0.008$ | - |
| | L | $0.25 \pm 0.008$ | $0.38 \pm 0.009$ | $0.30 \pm 0.007$ | $0.22 \pm 0.005$ | $0.18 \pm 0.005$ | $0.17 \pm 0.007$ | - |
| | $p$ | 0.0546 | 0.6709 | 0.6600 | 0.0678 | 0.3340 | 0.6634 | - |

**Figure 4:**
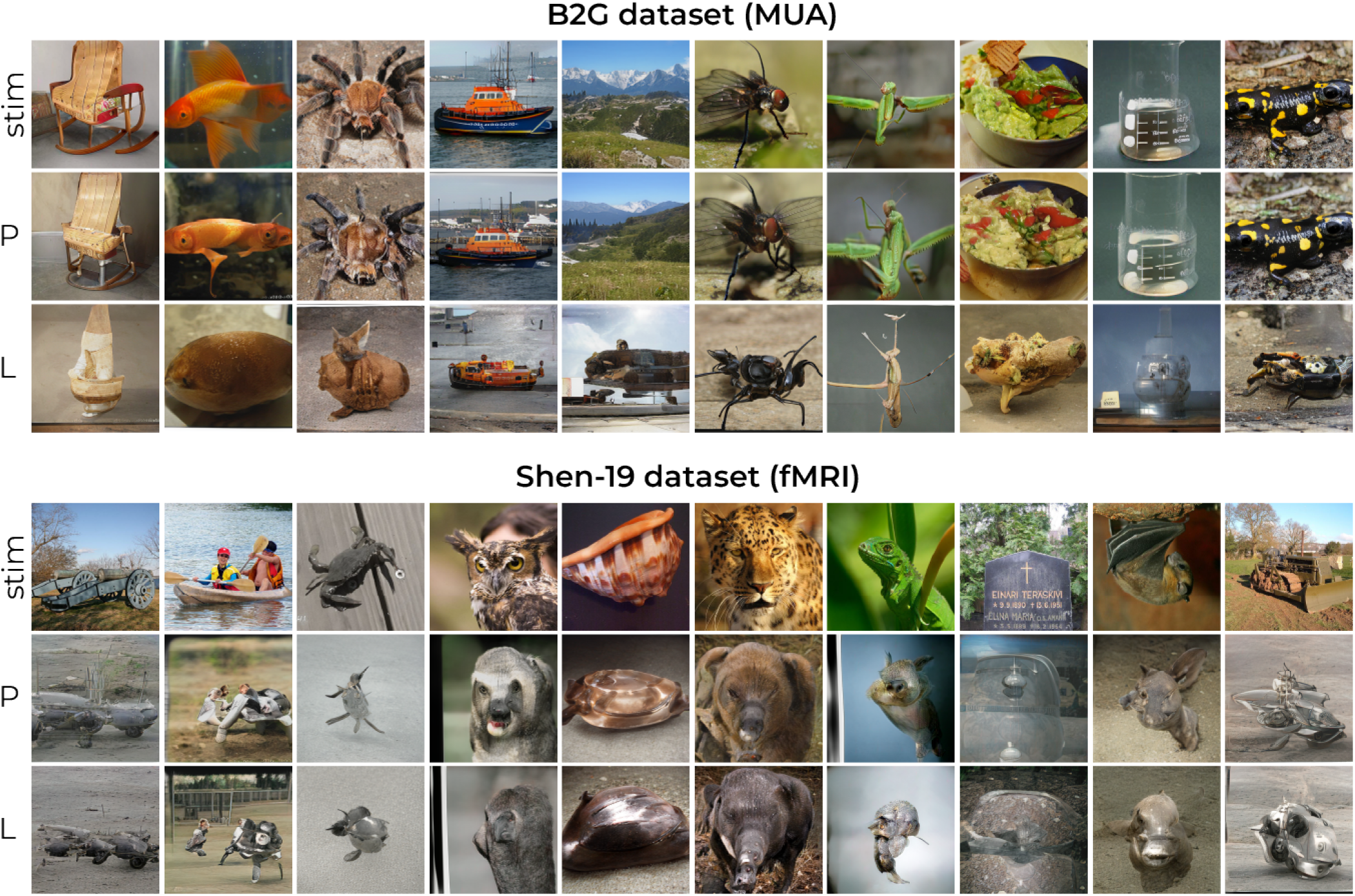
Reconstruction results. The upper and lower block show ten arbitrary yet representative examples from the B2G dataset (GAN-synthesized images) and Shen-19 dataset (natural photographs), respectively. The top rows display the stimuli, the middle rows the reconstructions by PAM (P) and the bottom rows the reconstructions by the linear decoder baseline (L).

The ability of PAM to perform on par with or better than the linear decoder confirms that it reconstructs stimuli with stateof-the-art fidelity. Unlike linear decoding, PAM offers a key interpretive advantage by providing insights into how attention is allocated across brain regions. Analysis of attention weights revealed that higher-order visual areas consistently received the most attention — specifically, IT in B2G and LOC in Shen-19 (Fig 5). In Shen-19, attention weights were more evenly distributed across visual areas, but there was still a gradual increase from early areas (V1-V4) toward LOC, accompanied by a notable dip at FFA. This pattern supports the idea that StyleGAN-XL’s *w*-latents predominantly encode high-level visual features, which correspond closely to activity in higherorder visual regions (Dado et al., 2024).

**Figure 5:**
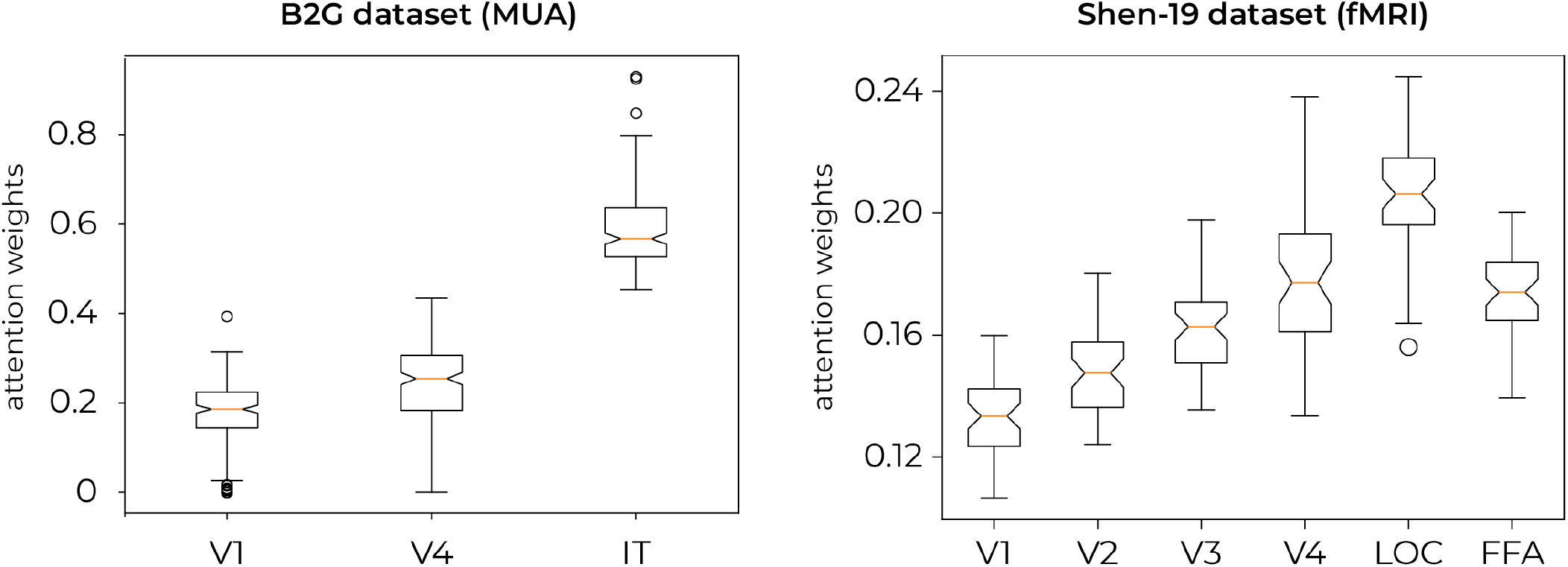
Distribution of the attention weights. The left panel illustrates the box plots of attention weights for the B2G dataset, derived from intracranial MUA recordings, across three regions of interest: V1, V4, and IT. The right panel displays attention weight distributions for the Shen-19 dataset, obtained from fMRI recordings, across six ROIs (V1, V2, V3, V4, LOC and FFA). Each box plot shows the median (orange line), interquartile range (box) and the full range excluding outliers (circles). For B2G, V1 received the lowest attention, while IT consistently received the highest, indicating a strong weighting toward higher-order visual processing. For Shen-19, attention weights were more evenly distributed across areas, but also showed a peak in the higher-order region LOC.

To break down decoding performance into its components, Fig 6 shows an example from B2G (**A**) and Shen-19 (**B**), illustrating how attention weights assign importance to the extracted values (i.e., the feature-rich transformations of neural activity that carry the stimulus information) and combine these into a single latent representation for reconstruction. Each brain area contributes different stimulus features, shaping the final decoded image. To see what specific stimulus properties each brain area is processing (Figs 6 and 7), we reconstructed these area-specific values by feeding them to the generator.

**Figure 6:**
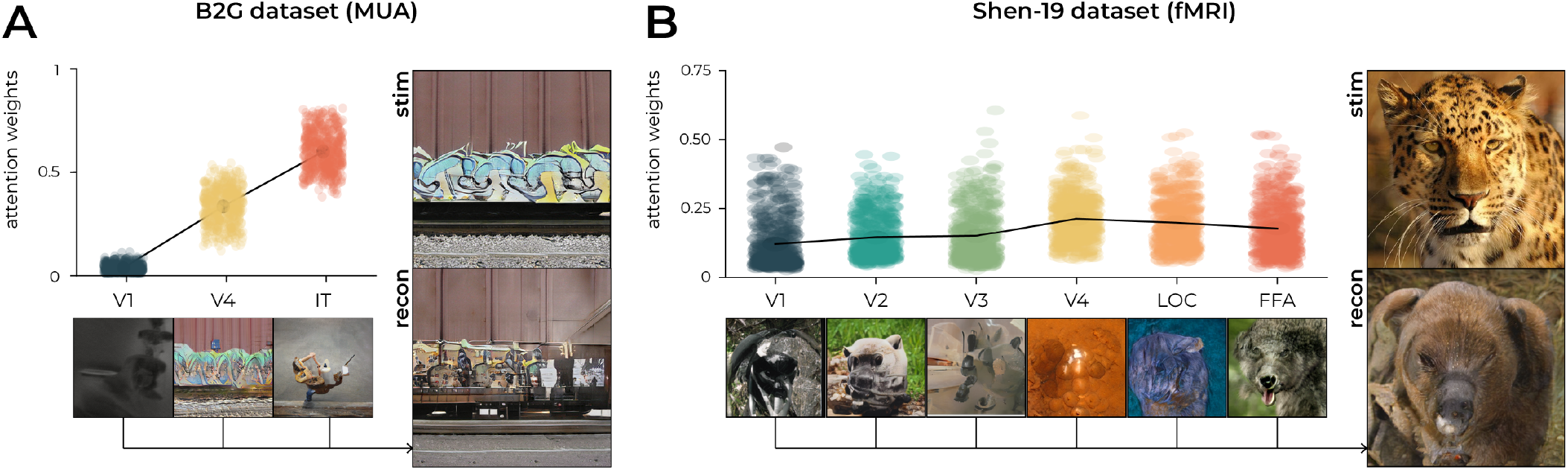
Attention-weighted values by PAM. The graphs visualize the distribution of 512-dimensional attention weights across the visual areas (V1, V4 and IT for B2G; V1, V2, V3, V4, LOC, and FFA for Shen-19) for two stimulus examples (‘stim’; on the right of the graph). The black lineplot denotes the mean attention per neural area. We can notice a gradual increase of attention from up- to downstream visual areas (more subtle for Shen-19). Below each label in the graph (*x*-axis), we visualized the visual information from the corresponding values by feeding them to the generator of the GAN. We then took a weighted combination of the values and the attention weights to obtain the final latent corresponding to the final reconstruction (‘recon’; displayed on the right, below the stimulus). For this example from B2G, particularly V4’s visualized value seems to resemble the stimulus. And also for the example from Shen-19, the warm colors and the dotted pattern from the panther seem to be reflected in the reconstructed value of V4 but not necessarily in the final reconstruction itself.

**Figure 7:**
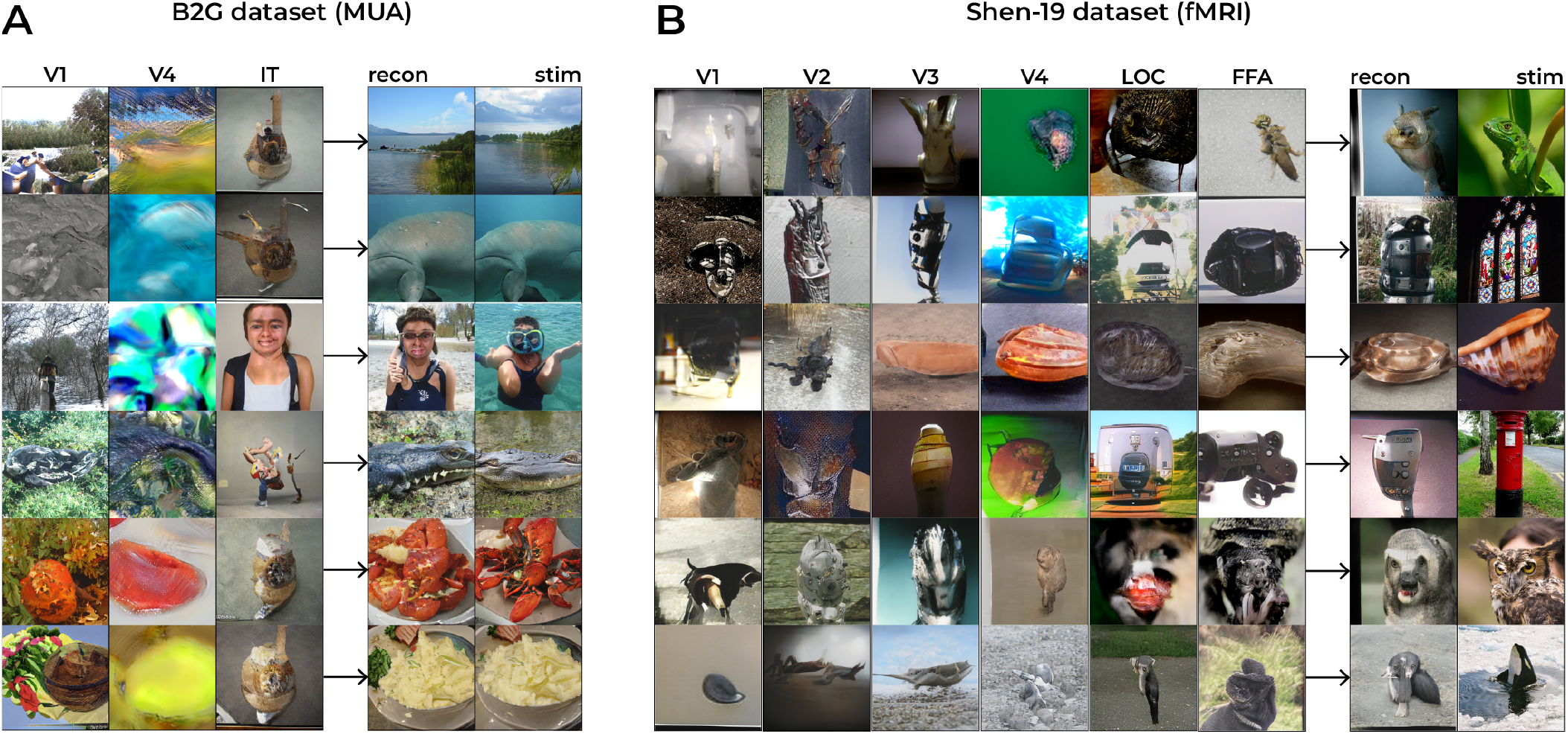
Reconstructed values. We visualized the information about the stimulus from each neural area by feeding their corresponding value to the generator of the GAN. For B2G, the reconstructed values from V1 seem to match the stimulus in basic outline, from V4 in color information and from IT in faces although the other reconstructions from this area seem rather meaningless despite the high assigned attention. Note that these stimuli are computer-generated such that the people in the third column do not really exist. For Shen-19, the reconstructed values from V1-2 seem to match the stimulus basic outlines as well, and V3 and V4 in shape and color information, respectively. The reconstructions from LOC and FFA seem to match the stimulus in terms of faces and contextual information.

- **B2G (MUA data)**. The reconstructed values from V1 primarily capture basic outlines, while those from V4 emphasize color information. In IT, the reconstructed values consistently reflected faces or animal-like features whenever the stimulus was animate. However, for inanimate stimuli, the IT values did not clearly represent meaningful stimulus features, despite IT receiving the highest attention weighting. This pattern was consistently observed across the entire test set—whenever a stimulus contained a face or an animal, its features were present in the IT-reconstructed values, but this was not the case for other object categories. Although the test set contained 200 images, we present only one example (i.e., the diver in Fig 7A) due to space constraints. Notably, IT received the strongest attention weighting, reinforcing its critical role in shaping the final latent representation, even when its direct reconstructions lacked clear details for inanimate stimuli.
- **Shen-19 (fMRI data)**. The reconstructed values from V1V2 primarily captured basic outlines, while V3 extracted more defined shapes. In V4, the values reflected color and texture information, whereas the higher-order areas (LOC and FFA) captured more contextual features. Integrating these values — weighted by attention — into a single latent vector produced meaningful reconstructions. Due to the limitations of fMRI data resolution, the final reconstructions were of lower quality compared to B2G, but they retained distinct stimulus characteristics, reflecting the contribution of different visual areas.

## Discussion

In this work, we introduced predictive attention mechanisms (PAMs), which learn task-driven output queries to prioritize relevant neural signals for reconstruction. By applying PAMs to neural decoding, we achieved state-of-the-art reconstructions while leveraging an interpretable attention mechanism to gain insight into the decoding process. Note that we do not claim PAM to be better than a conventional transformer, nor is it possible to make this comparison since it is not possible to derive output queries from neural data in the traditional sense.

Reconstructions from the B2G dataset (MUA data) were superior to those from Shen-19 (fMRI data). Intracranial MUA captures rapid, localized neural activity with high temporal and spatial resolution, providing high-quality signals. Because MUA allows for faster stimulus presentation than fMRI, it enabled the collection of a larger dataset. Combined with direct access to ground truth latent representations (StyleGAN-XL), this allowed for training a more complex model (five embedding blocks) without overfitting, making B2G an optimal dataset for reconstruction. In contrast, Shen-19 used fMRI, which measures slower, hemodynamic responses that indirectly reflect neural activity. This results in lower temporal resolution, greater noise susceptibility and a smaller dataset. Additionally, because real-world images lack pre-existing latents, we had to approximate them post-hoc, further compromising reconstruction accuracy. These factors limited model complexity — deeper embeddings led to overfitting, requiring a single-block embedding transformation. While MUA’s invasive nature limits its applicability, fMRI’s non-invasiveness comes at the cost of reduced precision, making reconstruction more challenging.

Our results showed a consistent trend of increased attention in downstream visual areas, though the specific allocation of attention varied across individual examples. This demonstrates how PAM does not apply a fixed attention pattern but instead dynamically adjusts its weighting based on how well neural responses match the learned queries. In other words, PAM adapts its focus to the unique neural characteristics of each stimulus, emphasizing regions that provide the most relevant information for reconstruction. However, while PAMs enhance interpretability by providing attention maps, the rationale behind specific weight assignments remains opaque, making it challenging to fully disentangle the underlying mechanisms driving its performance. For Shen-19, attention progressively increased across higher-order visual areas but dipped at the FFA, despite still exceeding early visual areas (V1-V3). This could be attributed to the dataset’s stimulus composition: while it included animal faces (e.g., a panther and an owl) and human figures engaged in activities (e.g., a person playing the harp and two people in a canoe), it lacked close-up human faces — typical triggers for strong FFA activation. The absence of such stimuli could explain the reduced focus on FFA.

Visualizing the values associated with each brain region provides insights into what stimulus information was decoded from different areas. This is because these values reflect the specific features extracted at each stage of the visual hierarchy. We found that V2 and V3 primarily captured basic outlines and shapes, while V4 represented colors and textures. More downstream areas encoded higher-order attributes (e.g., object identities, contextual relationships) that integrate basic sensory information. Interestingly, although PAM allocated greater attention to downstream areas, their reconstructed values often showed less direct visual similarity to the original stimuli than those from earlier regions (e.g., V4). However, when all values were integrated using attention weights, the final reconstructions remained highly accurate. This suggests that deeper areas may act as directional vectors in latent space, guiding the model toward the correct overall perceptual reconstruction. So, even if their value reconstructions appear less visually aligned with the stimulus, these areas play a crucial role in shaping the final output. By assigning greater weight to their decoded values, PAM utilizes the richer semantic and contextual representations processed in higher-order visual regions to refine the final output.

Future work should investigate how PAM dynamically shifts attention depending on the level of visual representation being decoded. In our study, GAN-based decoding aligned neural responses with a structured latent space that best captured high-level visual features, resulting in increased attention to higher-order visual areas. Conversely, decoding lowlevel features (e.g., Gabor-like edge representations) would likely shift attention toward early visual areas, where these features are processed. Understanding how PAM allocates attention across the visual hierarchy could reveal how different brain regions contribute to perception at varying levels of abstraction. Leveraging PAM to track attention shifts in realtime would offer a new way to investigate hierarchical information flow in the brain. Ultimately, integrating different levels of representations within a unified decoding framework could further improve neural reconstructions. This may contribute to advances in brain-computer interfaces (BCIs) and neuroprosthetics, particularly in sensory processing. Understanding how attention is distributed across brain areas could also inform clinical interventions, such as targeted neural stimulation for visual disorders.

Neural decoding research raises ethical concerns, particularly regarding mental privacy and the potential misuse of technology. However, our models rely on carefully controlled datasets and require full subject cooperation, meaning they cannot be applied to individuals outside the training set. They reconstruct only externally perceived images and do not decode internal thoughts, mental imagery or dreams.

Beyond neural decoding, PAMs hold promise for broader applications. Future work should explore their integration across different domains and modalities.

## Acknowledgements

This work was supported by the DBI2 project through the NWO Gravitation programme (grant number 024.003.013).

